# Fluorescence Correlation and Cross-Correlation Spectroscopy Unveil Cytoplasmic mRNP Composition and Dynamics

**DOI:** 10.1101/2020.07.20.212308

**Authors:** Àngels Mateu-Regué, Jan Christiansen, Christian Hellriegel, Finn Cilius Nielsen

**Author notes:** Correspondence: Finn Cilius Nielsen, Center for Genomic Medicine Rigshospitalet, Copenhagen, Denmark, E.mail.

## Abstract

Understanding the mRNA life cycle requires analysis of the dynamic macromolecular composition and stoichiometry of mRNPs. Fluorescence correlation and cross-correlation spectroscopy (FCS and FCCS) are appealing technologies to study mRNP complexes because they readily provide information about the molecular composition, stoichiometry, heterogeneity and dynamics of the particles. We developed FCS protocols for analysis of live cells and cellular lysates, and demonstrate the feasibility of analysing common cytoplasmic mRNPs composed of core factor YBX1, IMPs (or IGF2BPs) and their interactions with other RNA binding proteins such as PABPC1, ELAVL2 (HuB), STAU1 and FMRP. FCCS corroborated previously reported RNA dependent interactions between the factors and provided an estimate of the relative overlap between the factors in the mRNPs. In this way FCS and FCCS provide a new and useful approach for the quantitative and dynamic analysis of mRNP macromolecular complexes that may complement current biochemical approaches.

## INTRODUCTION

The mRNA life cycle begins in the nucleus where the newly transcribed precursor mRNA undergoes splicing, capping and polyadenylation. The mature mRNA subsequently exits the nucleus through the nuclear pore after which it becomes transported, translated and/or subsequently degraded. All steps involve the sequential attachment of RNA-binding proteins governing the individual processes (1). In the cytoplasm, individual transcripts are coated with different RNA-binding proteins, and the small messenger ribonucleoprotein particles (mRNPs) represent the cellular transcriptome (2). Deficiencies in the attached RNA-binding proteins can lead to deficiencies in cell functionality and ultimately to disease (3–8).

A deeper understanding of the post transcriptional control of gene expression and its role in pathological processes requires capability to examine the molecular composition of particular mRNPs. mRNP analysis has to a large extent relied on biochemical analysis of isolated mRNP - employing antibodies or tagged baits in pull-down analysis or fluorescence sorting in combination with western analysis and/or mass spectrometry (9–13). Although these methods have provided an impressive insight into the components of particles, they are incompatible with an analysis of live cells and have a number of limitations with respect to stoichiometry, heterogeneity and subcellular resolution. Fluorescence Correlation and Cross-Correlation Spectroscopy (FCS and FCCS, respectively) are appealing alternatives for the analysis of protein interactions and the composition of complex protein assemblies because they can be applied to live cells and cellular lysates. By recording the fluorescence fluctuations produced by labelled molecules or proteins entering and leaving a small focal volume over time and their subsequent analysis by autocorrelation, FCS instantly provides information about the diffusion time, concentration and aggregation of fluorescent molecules (14) (Figure 1A). FCCS is a variant of FCS and is based on a two-fluorophore analysis. By analysing the cross-correlation between molecules or proteins labelled with two spectrally distinct fluorophores it is moreover possible to establish if two factors associate or are part of the same macromolecular complex (15).

**Figure 1.**
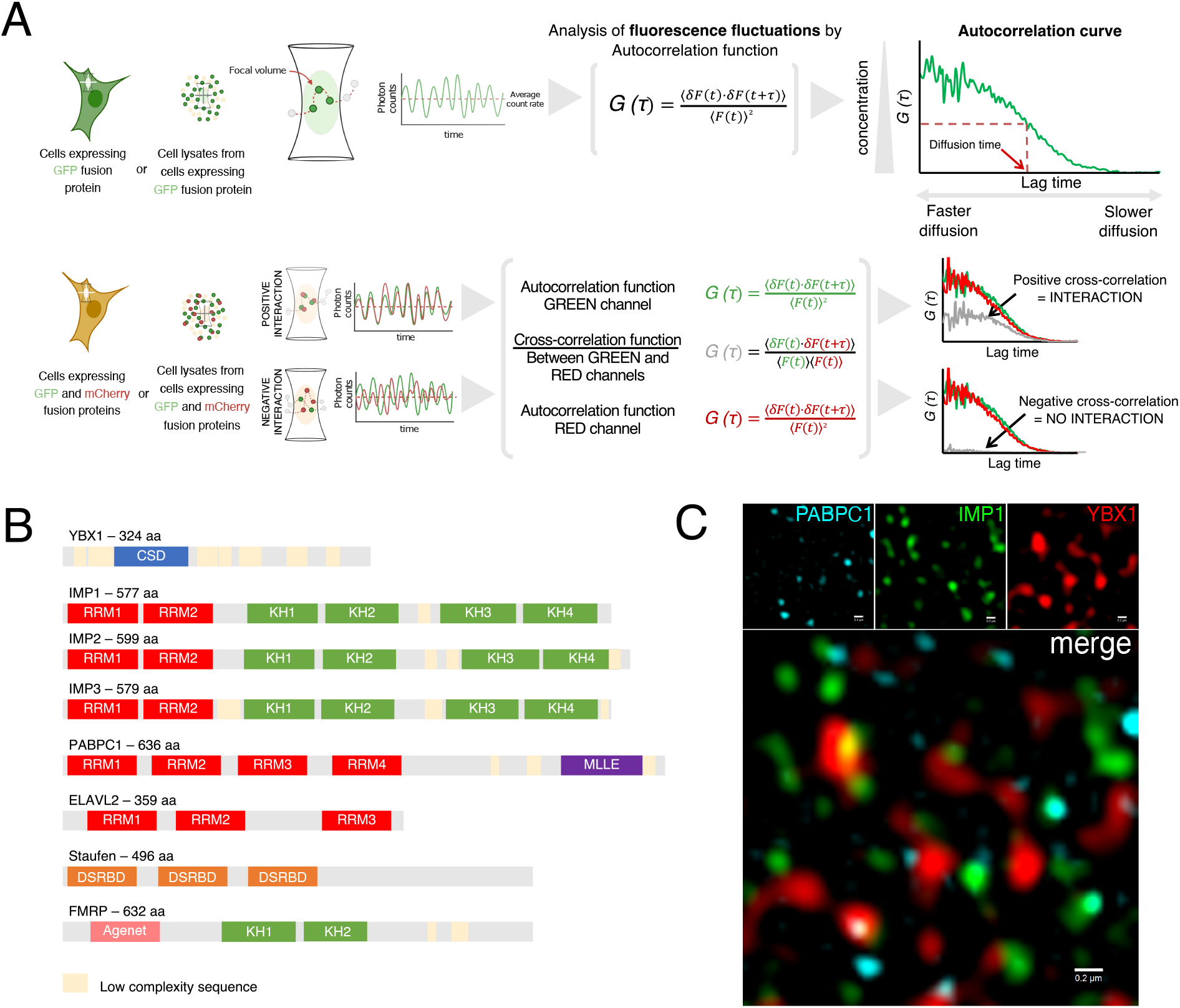
Overview of Fluorescence Correlation Spectroscopy (FCS) and Fluorescence Cross-correlation Spectroscopy (FCCS) methods and RNA-binding proteins used in the study. **A.** FCS measurements are performed either in cells expressing a protein of interest fused to a GFP tag or in cell lysates from cells expressing the GFP fusion protein. Focal volume is positioned in a specific cellular location (live cell FCS) or in a fluorescent protein solution (cell lysate FCS). Fluorescence fluctuations are recorded and are subsequently analyzed by the autocorrelation function, which result in the autocorrelation curve, used to determine the diffusion time of the examined GFP-tagged protein. FCCS is based on the combined FCS measurements of two spectrally-distinct fluorescent proteins i.e. GFP and mCherry. Autocorrelation function is used for analyzing diffusion of GFP- and mCherry-tagged proteins and correlation function is also applied between channels (cross-correlation). Interacting GFP- and mCherry-tagged diffuse synchronously through the focal volume, therefore cross-correlation curve (grey) is positive. In contrast, non-interacting GFP- and mCherry-tagged proteins diffuse independently from each other yielding a flat or negative cross-correlation curve. **B.** Schematic representation of the different proteins used in this assay with their RNA binding domains and predicted low complexity sequences (CSD: Cold-shock domain; RRM: RNA recognition motif, KH: K-homology, DSRBD: Double-stranded RNA-binding domain). **C.** Structured Illumination Microscopy (SIM) image of HeLa cell cytoplasmic mRNPs obtained by indirect immunofluorescence staining of their major protein components: YBX1 (Alexa Fluor 647, red), IMP1 (Alexa Fluor 568, green) and PABPC1 (Alexa Fluor 488, cyan). Scale bar = 0.2 μm

We employed FCS and FCCS for characterization of cytoplasmic mRNPs examining associations of a number of well established cytoplasmic core mRNA binding proteins, including the mRNP core proteins YBX1, IMP1 and PABPC1 as well as IMP2, IMP3, ELAVL2 (HuB), STAU1 and FMRP (1,4,16–18) (Figure 1B and 1C). FCS and FCCS are fast and reproducible and data obtained in live cells is reconciled in cell lysates. We moreover show how FCS can be used to study the stoichiometry of mRNP complexes. FCCS corroborated previously reported RNA dependent interactions of the factors and complemented the interaction data with insights about the likelihood of the interaction and the heterogeneity. We infer that FCS and FCCS provide a valuable approach for the analysis of mRNP.

## RESULTS

### FCS of mRNPs in live cells and cell lysates

Fluorescence intensity fluctuations in HeLa cells expressing GFP-tagged YBX1, IMP1 and IMP1_KH1-4mut_, an IMP1 mutant with impaired RNA-binding by mutation of the GXXG loops in the 4 KH domains of IMP1 (19) (Figure 2A), were recorded in the cytoplasm of live cells and in cell lysates (Figure 2C-F). GFP was included as a reference in both live cells and cell lysates (Figure 2F). Cells expressing GFP-tagged proteins were localized by confocal imaging and the focal volume was positioned at arbitrary locations in the cytoplasm as shown in Figure 2A. Cell lysates were prepared by the addition of iso-osmolar lysis buffer containing non-ionic detergent to cells expressing GFP-tagged proteins, in order to release the cytoplasmic content. In the case of cell lysates, localization of the focal volume was not needed since the fluorescent molecules or complexes were not confined in a specific place but were diffusing homogeneously in the lysate solution. Live cell FCS measurements were carried out in different cells from a single transfection and also from different transfections and cell lysate FCS measurements were recorded from different cell transfections. We found the measurements highly reproducible (Supplementary Figure 2). Therefore, representative measurements of each GFP-tagged factors are shown in Figure 2.

**Figure 2.**
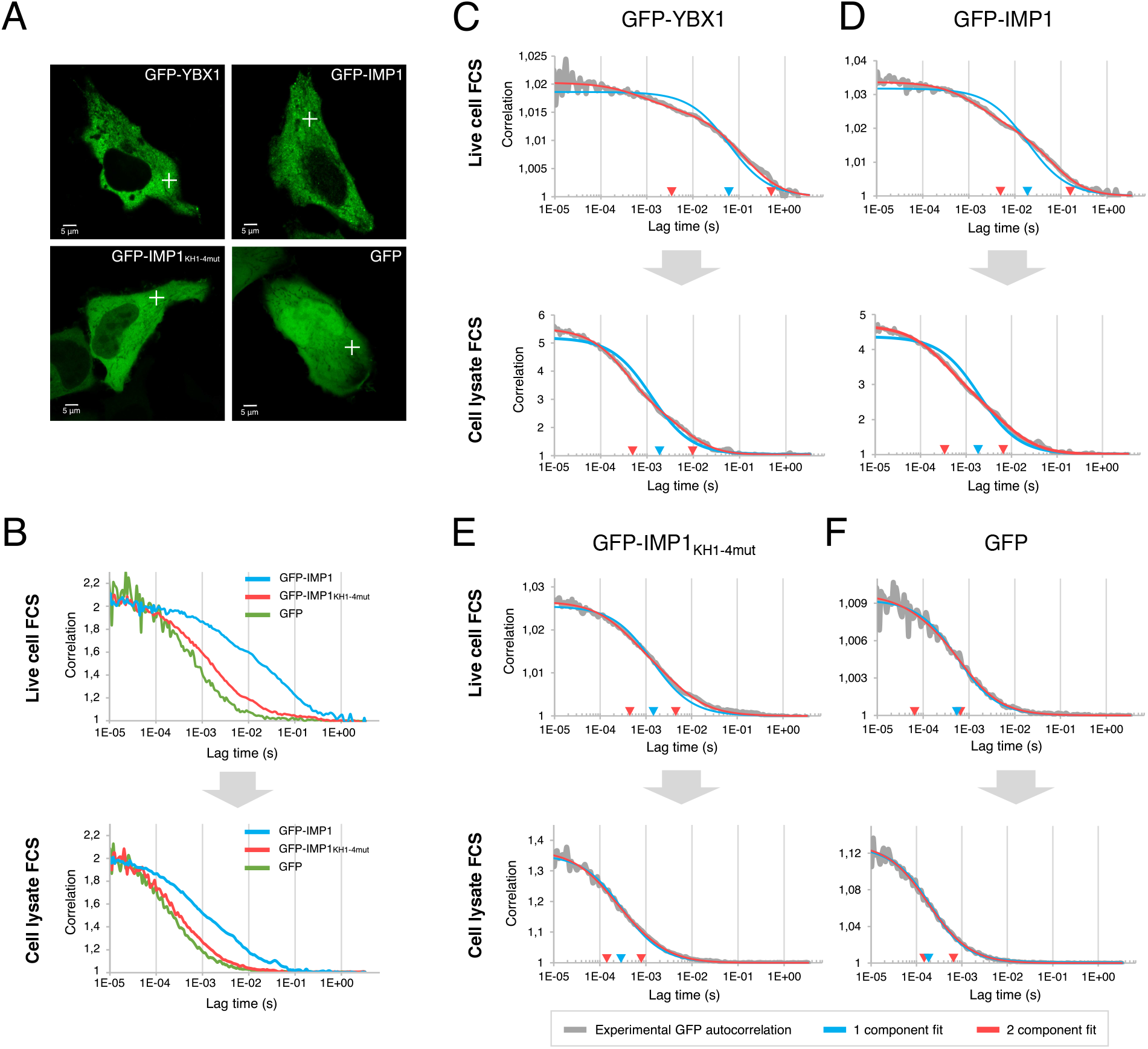
Comparison of live cell and cell lysate Fluorescence Correlation Spectroscopy (FCS). **A.** Confocal microscopy images of HeLa cells expressing GFP-YBX1, GFP-IMP1, GFP-IMP1_KH1-4mut_ (RNA-binding impaired mutant) and GFP. Crosses illustrate how arbitrary points in the cytoplasm were chosen to perform live cell FCS. Scale bar = 5 μm. **B.** Autocorrelation curves (normalized from 1 to 2) from GFP-IMP1 (mRNP), GFP-IMP1_KH1-4mut_ and GFP measurements in live cells (top) and cell lysates (bottom). **C.** GFP-YBX1, **D.** GFP-IMP1, **E.** GFP-IMP1_KH1-4mut_ and E. GFP representative autocorrelation curves from live cells (top) and cell lysates (bottom). Fittings to 1-component (blue) and 2-component (red) models are displayed in the graphs together with the experimental autocorrelation curves. Blue and red arrows in x axis (lag time, sec) represent the diffusion coefficients corresponding to the fit to 1-component (blue) and 2-component (red) diffusion models.

In comparison to live cell diffusion, the relative distribution of the autocorrelation curves of GFP, GFP-IMP1_KH1-4mut_ and GFP-IMP1 was kept in cell lysates, but in agreement with the lower viscosity and unrestrained motions in lysates, diffusion was in general faster. The diffusion constant of the mRNP component GFP-IMP1 was slower than GFP, in accordance with the observed embedment in the mRNP complex. In agreement with an RNA dependent association of GFP-IMP1 with mRNPs, disruption of its RNA-binding ability increased its diffusion almost to the level of free GFP (Figure 2B). The average difference between live cells and lysates was quantified and compared by fitting the experimental autocorrelation curves to a single-component diffusion model (Figure 2C-F, blue line and arrows), and this demonstrates that diffusion is about one order of magnitude slower in live cells than cell lysates. However, the GFP-YBX1 and GFP-IMP1 autocorrelation curves did not fit with a 1-component diffusion model, indicating the presence of an additional dynamic process. We subsequently added complexity to the model with a 2nd component, which improved the experimental fitting (Figure 2C and 2D, red line and arrows). Fittings in both live cells and lysates for GFP-IMP1 and GFP-YBX1 were improved when adding a second component to the model, indicating that the cell lysates reconcile the situation in live cells. GFP-IMP1_KH1-4mut_ (Figure 2E) fitted better to a 1-component diffusion model than GFP-YBX1 and GFP-IMP1, but the fit was still significantly better with a 2-component model. This is probably due to a residual RNA-binding activity of the protein. In the case of GFP (Figure 2F), the single component model in both conditions is sufficient, a fact that reconciles the unrestrained 3D diffusion of this protein in live cells as well in cell lysates. Taken together, we conclude that FCS is able to depict the motion of macromolecular mRNP assemblies and that analysis of lysates largely recapitulates the *in vivo* conditions and at the same time improves accuracy, reproducibility and simplicity.

### mRNP stoichiometry

mRNPs are composed of different RBPs, and mRNAs exhibit several binding sites for a particular RBP. We examined the stoichiometry of IMPs, which bind to single-stranded CA-rich stretches, so a particular mRNA may in principle harbour one or two handfuls of IMPs depending on the size of the transcript. Cells expressing GFP-IMP1, GFP-IMP2, GFP-IMP3 and GFP-IMP1_KH1-4mut_ were first lysed in iso-osmolar lysis buffer and cell lysates were subjected to FCS (Figure 3A, top row). From the recorded fluorescence intensity fluctuations, mRNPs containing multiple GFP-tagged IMPs can be observed directly as peaks when the count rate (kHz) is plotted as a function of time. The variation of the number of associated IMP molecules is reflected by the different count rates of the individual peaks. In contrast, peaks were not observed in the recorded fluctuations for GFP-IMP1_KH1-4mut_ because the mutant exist as a monomer. The same pattern is observed for GFP-IMP1, GFP-IMP2 and GFP-IMP3 after treatment with RNase A (Figure 3A, middle row) or lysis in a hypertonic lysis buffer, which disrupts RNA-protein interactions (Figure 3A, bottom row).

**Figure 3.**
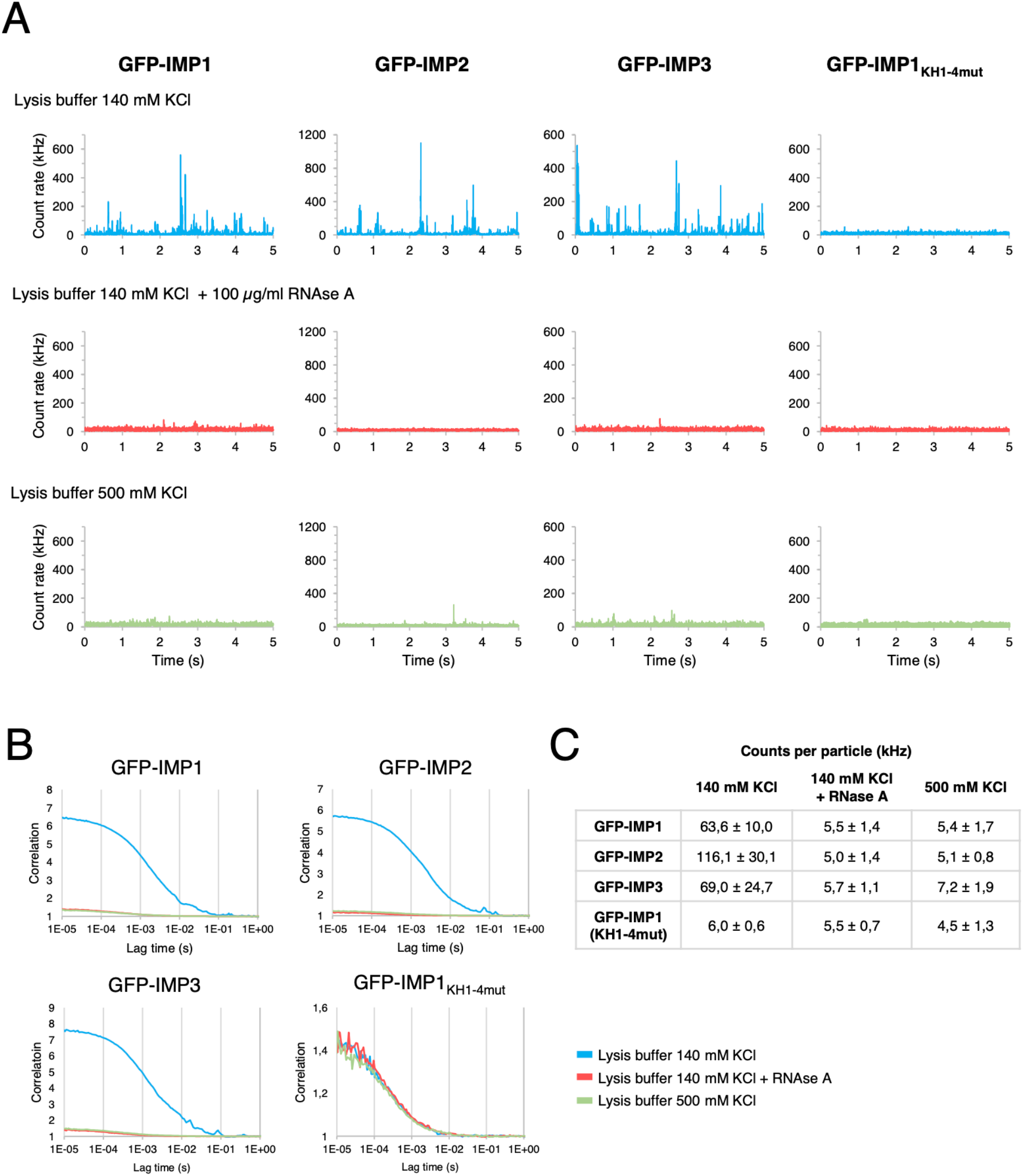
IMP protein stoichiometry in mRNPs. **A.** Fluorescence intensity measurements (kHz) in time (5 seconds) of cell lysates from HeLa cells transfected with GFP-IMP1, GFP-IMP2, GFP-IMP3 and GFP-IMP1_KH1-4mut_. Cells were lysed using iso-osmolar lysis buffer (blue, top row), subsequently treated with 100 μg/mL RNase A (red, medium) or lysed with hyperosmolar lysis buffer in order to break protein-RNA interactions (green, bottom). **B.** Fluorescence autocorrelation curves of HeLa cell lysates transfected with GFP-IMP1, GFP-IMP2, GFP-IMP3 or GFP-IMP1_KH1-4mut_ under different buffer conditions: iso-osmolar lysis buffer (140 mM KCl), iso-osmolar lysis buffer (140 mM KCl) + RNAse A treatment and hyperosmolar lysis buffer (500 mM KCl). **C.**Counts per particle (CPP) averages ± standard deviations of n=6 biological replicates in cell lysates.

The amplitude of the autocorrelation curve is inversely correlated to the concentration of the fluorescent molecules or particles in solution, so highly concentrated fluorescent particles will yield lower amplitudes and vice versa. As shown in Figure 3B, when IMPs are part of mRNPs, multiple IMPs are bound to the same mRNA. Therefore the concentration of fluorescent particles is lower but the brightness of each particle is higher because it contains several fluorophores. After treatment with RNase A or lysis in hypertonic lysis buffer, IMPs dissociate from mRNPs and exist as monomers. Consequently, the concentration of the fluorescent species is higher whereas brightness is lower. The amplitude of GFP-IMP1_KH1-4mut_ remained unchanged in all conditions because this protein exists as a monomer in all conditions (Figure 3B).

By dividing the total fluorescence (total count rate) with the number of molecules in the focal volume (derived from the amplitude of the autocorrelation curve) FCS allows determination of the number of fluorescent molecules or “labels” present in a given complex (counts per particle or CPP, or sometimes referred to as counts per molecule CPM). To determine the number of IMPs in each mRNP, we compared the counts per particle in cell lysates of cells transfected with GFP-IMP1, GFP-IMP2 or GFP-IMP3 with GFP-IMP1_KH1-4mut_. Moreover, lysates were subjected to RNase A treatment or lysed with a hypertonic buffer to obtain the monomeric brightness. Comparing the mRNP brightness with the brightness of the monomer (GFP-IMP1_KH1-4mut_), the analysis shows that mRNPs on average contain 10 GFP-IMP1, 19 GFP-IMP2 and 11 GFP-IMP3 molecules (Figure 3C). We noticed that even though the same amount of plasmid was transfected, GFP-IMP2 was expressed at higher levels (~2 fold) compared to GFP-IMP1 and GFP-IMP3 (Supplementary Figure 3). This could be due to an increased IMP2 half-live in live cells. When CPP were normalized by the expression level of each protein, we could see that no differences for GFP-IMP1, GFP-IMP2 and GFP-IMP3 were observed (Supplementary Figure 3). Consequently, we infer that all IMP members have the same ability to bind to mRNA and that their expression level determines the relative abundance of IMPs in individual mRNPs. The results moreover imply that the expressed RBPs are within the biological range since the mRNPs are able to accommodate more IMPs.

### mRNP core protein interactions

Once we established the behaviour of the mRNP components by FCS, we subsequently explored the feasibility of Fluorescence Cross-Correlation Spectroscopy (FCCS) to depict associations between different RNA-binding proteins. We initially tested the interactions between YBX1 and IMP1 or PABPC1, that previously have been demonstrated to constitute the major elements of cytoplasmic mRNPs (1). Cytoplasmic YBX1 coats mRNAs along their entire length by attachment to the sugar-phosphate backbone (17,20), whereas IMPs have a preponderance for loops and 3’UTRs (21).

HeLa cells were co-transfected with mCherry-YBX1 and GFP-IMP1 and the reverse pair mCherry-IMP1 and GFP-YBX1. PABPC1-GFP was co-transfected with mCherry-YBX1. As shown previously in Figure 2, cell lysates prepared by non-ionic iso-osmolar detergent solubilisation provide a good representation of the situation in live cells. By analysing the total cellular content when performing cell lysis, measurements are more homogeneous since the complexity is reduced when compared to the situation in a live cell. Moreover, a lower concentration of the tagged complexes or proteins yields better FCS measurements, due to the higher amplitude. In addition, differences or bias in measurements due to e.g. cytoplasmic localization can be avoided.

As a positive control for cross-correlation, a mCherry-GFP fusion protein emitting in both green and red channels was employed, whereas the negative control consisted of a co-transfection of mCherry and GFP in separate plasmids. As shown in Figure 4A, positive cross-correlation is observed with mCherry-GFP fusion protein while cross-correlation is not observed when fluorescent proteins are expressed as separate proteins. Due to photobleaching, misfolding or fluorescent proteins being “off” or in dark states (22), the cross-correlation is not expected to be 100%, as illustrated by the mCherry-GFP fusion protein.

**Figure 4.**
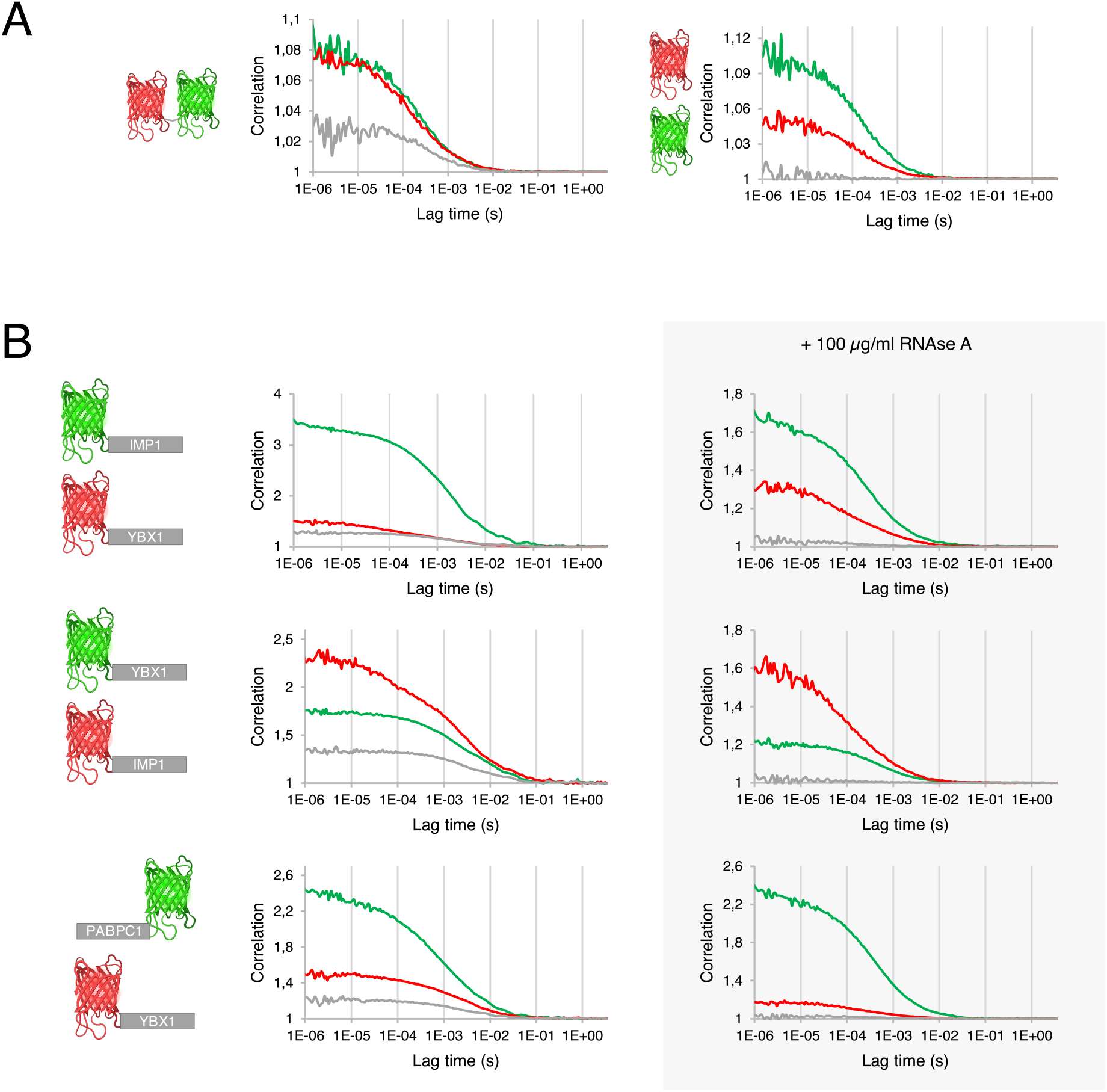
Protein-protein interaction quantification in cell lysates between three major mRNP components: YBX1, IMP1 and PABPC1. **A.** Cell lysate Fluorescence Cross-correlation Spectroscopy (FCCS) measurements of cells expressing a mCherry-GFP fusion protein (positive control for cross-correlation, left) and mCherry/GFP transfected in separate plasmids (negative control for cross-correlation, right). **B.** Cell lysate FCCS measurements of cells expressing GFP-IMP1 and mCherry-YBX1 (top), GFP-YBX1 and mCherry-IMP1 (middle) and PABPC1-GFP and mCherry-YBX1 (bottom). RNase A was added to the lysates and changes in cross-correlation were measured (right graphs).

mCherry-YBX1 exhibits a positive cross-correlation with both GFP-IMP1 and PABPC1-GFP. Cross-correlation is very high with GFP-IMP1, meaning that GFP-IMP1 containing mRNPs encompass mCherry-YBX1 to a great extent (Figure 4B, top). At the same time, not all GFP-YBX1 containing mRNPs have have mCherry-IMP1 in them, as observed in a positive but lower cross-correlation with GFP-YBX1. This confirms YBX1 as a general and universal factor of mRNPs and IMP1 as a factor binding to a subset of mRNAs (Figure 4B, middle). Positive cross-correlation between PABPC1-GFP and mCherry-YBX1 indicates interaction between the factors but in agreement with the widespread and general distribution of PABPC1, to a lesser extent. Finally, cross-correlation was in all of the cases lost when the cell lysates were treated with RNase A, demonstrating that the interactions between IMP1-YBX1 and PABPC1-YBX1 are RNA dependent.

### Coexistence of IMP1 and YBX1 with ELAVL2, STAU1 and FMRP in mRNPs

Different mRNA binding proteins have been described to interact by immunoprecipitation analysis but whether these mRNP complexes are homogeneous or heterogeneous, as well as the likelihood of the interaction between the RNA-binding proteins, is unresolved. To address the likelihood of the interaction between IMP1 or YBX1 – one abundant and one core component of mRNPs, respectively – was assessed by FCCS. Cells were transfected with mCherry-IMP1 in combination with GFP-tagged IMPs, ELAVL2, STAU1 or FMRP and fluorescence measurements were recorded in cell lysates (Figure 5A). We calculated the average cross-correlation/mCherry-IMP1 autocorrelation ratio across six biological replicates by dividing the cross-correlation amplitude with the mCherry-IMP1 amplitude (Figure 5C). mCherry-IMP1 cross-correlates to a high degree with GFP-IMP1, GFP-IMP2 and GFP-IMP3, showing that these proteins are part of the same mRNP complexes. The cross-correlation ratio for IMP1 tagged with two different fluorescent proteins is close to the maximum cross-correlation, and it is even higher than 50%. This is probably due to the presence of multiple IMP molecules in the mRNP, which makes the occurrence of completely “off” states less likely. GFP-ELAVL2 shows an intermediate degree of cross-correlation with mCherry-IMP1, meaning that ELAVL2 is present in mCherry-IMP1 containing mRNPs but to a lesser extent. On the other side, cross-correlation with GFP-STAU1 or GFP-FMRP and mCherry-IMP1 is very low, indicating that the proteins associate significantly less with mCherry-IMP1 containing mRNPs, in contrast to IMP1, IMP2, IMP3 and ELAVL2. Moreover, the interaction of all the tested GFP-tagged factors is RNA dependent, since cross-correlation is lost when the cell lysate is treated with RNase A (Figure 5C). The results obtained by FCCS were compared to immunoprecipitation of 3xFLAG-IMP1 from TREX 293 3xFLAG-IMP1 cells. As shown in Figure 5B, GFP-IMP1, GFP-IMP2, GFP-IMP3 and GFP-ELAVL2 can be immunoprecipitated with 3xFLAG-IMP1 while GFP-STAU1, GFP-FMRP and GFP were not detected in the immunoprecipitate. Taken together, we infer that FCCS measurements are in agreement with immunoprecipitation data although FCCS appears to be more sensitive.

**Figure 5.**
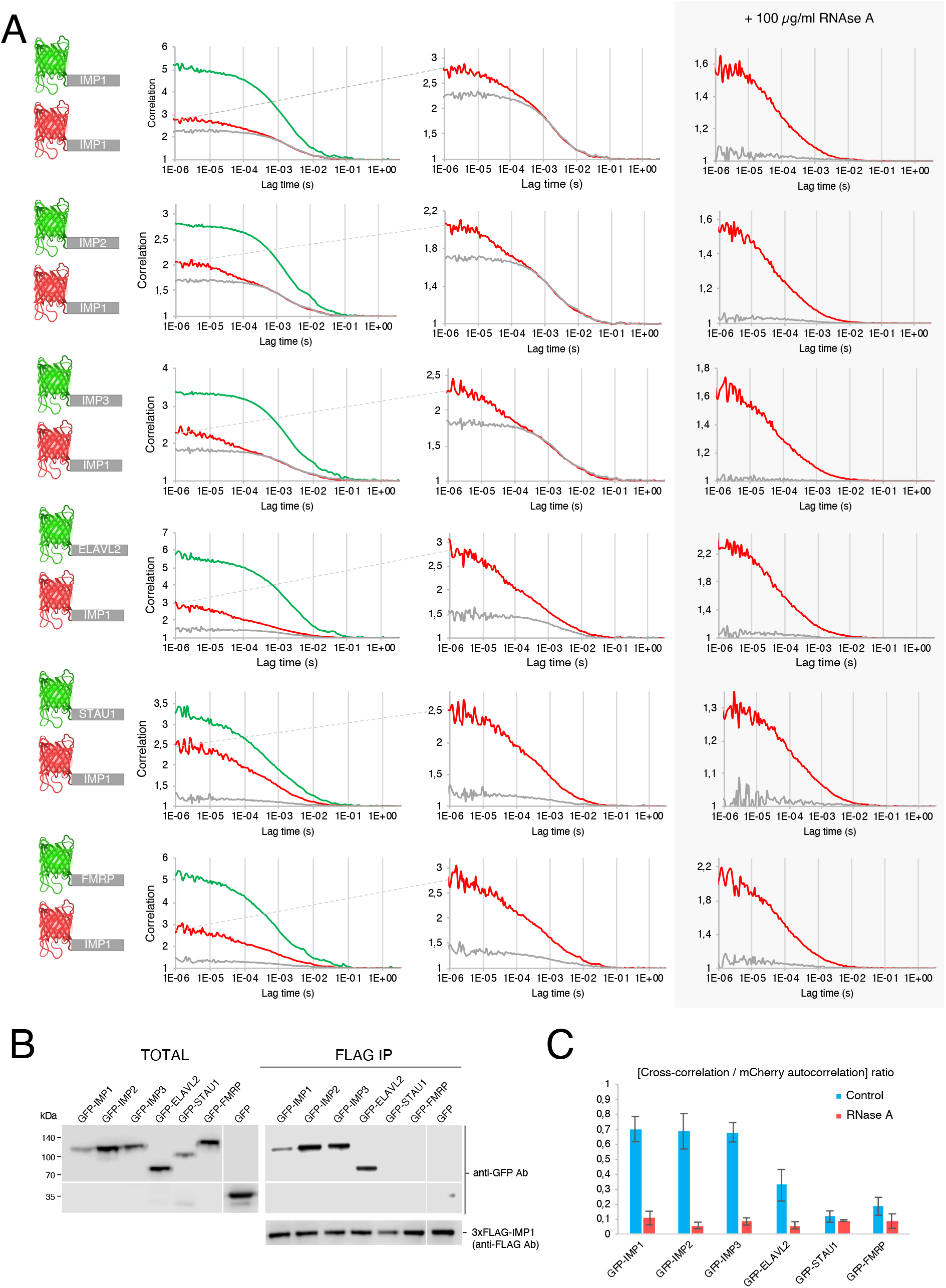
Protein-protein interaction quantification in cell lysates between mCherry-IMP1 and GFP-tagged IMP2, IMP3, ELAVL2, STAU1 and FMRP proteins. **A.** Representative Fluorescence Cross-correlation Spectroscopy measurements of HeLa cell lysates co-transfected with mCherry-IMP1 and GFP-IMP3, GFP-IMP2, GFP-ELAVL2, GFP-STAU1 or GFP-FMRP (left). Relative cross-correlation in respect to mCherry-IMP1 before (middle) or after treatment with 100 μg/mL RNase A (right). **B.** Western Blot of total lysate (left) and FLAG immunoprecipitated (right) fractions of TREX 293 3xFLAG-IMP1 cells transiently transfected with GFP-IMP1, GFP-IMP2, GFP-IMP3, GFP-ELAVL2, GFP-STAU1, GFP-FMRP and GFP (negative control). **C.** Average cross-correlation / mCherry-IMP1 autocorrelation ratios (± SD) showing different degrees of interaction depending on the GFP-tagged protein across n=6 biological replicates.

We subsequently performed FCCS for the same GFP-tagged proteins but this time employing mCherry-YBX1 (Figure 6A). Similar to mCherry-IMP1, GFP-tagged IMPs and ELAVL2 presented a high cross-correlation with mCherry-YBX1 mRNPs while STAU1 and FMRP cross-correlation was significantly lower (Figure 6B). However, cross-correlation of IMPs is not as high with mCherry-YBX1 as with mCherry-IMP1, and cross-correlation with GFP-ELAVL2 is slightly higher than with mCherry-IMP1. The relative overlaps with IMP1 and YBX1 are summarized in Supplementary Figure 4.

**Figure 6.**
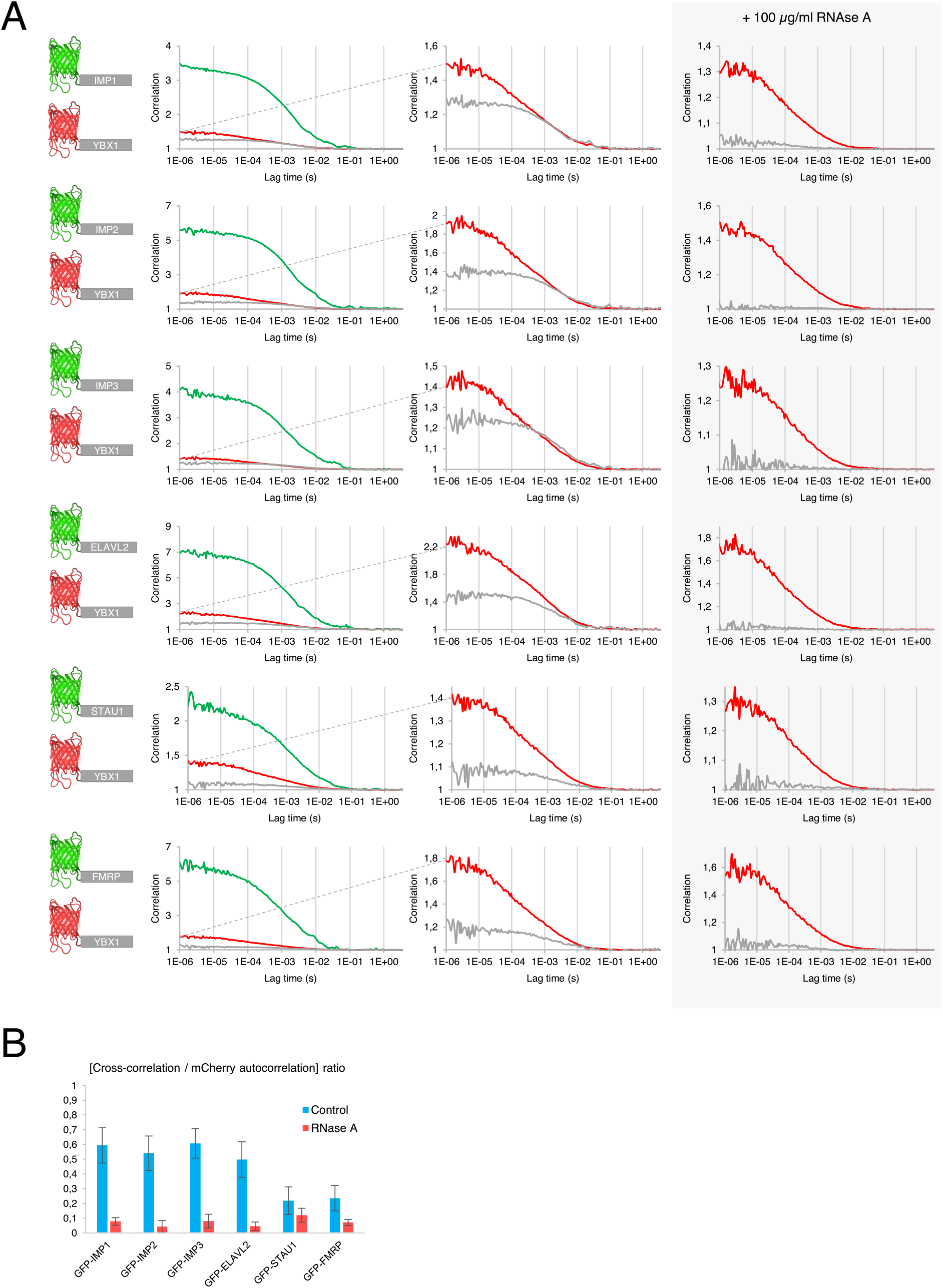
Protein-protein interaction quantification in cell lysates between mCherry-YBX1 and GFP-tagged IMP2, IMP3, ELAVL2, STAU1 and FMRP proteins. **A.** Representative Fluorescence Cross-correlation Spectroscopy measurements of HeLa cell lysates co-transfected with mCherry-YBX1 and GFP-IMP2, GFP-IMP3, GFP-ELAVL2, GFP-STAU1 or GFP-FMRP (left). Relative cross-correlation in respect to mCherry-YBX1 before (middle) or after treatment with 100 μg/mL RNase A (right). **B.** Average cross-correlation / mCherry-YBX1 autocorrelation ratios (± SD) showing different degrees of interaction depending on the GFP-tagged protein across n=6 biological replicates.

Finally, FCCS with mCherry-YBX1 was performed in live cells (Figure 7) to compare the above described associations. In agreement with the data shown in Figure 2, the autocorrelation curves shift approximately an order of magnitude. With the exception of GFP-FMRP, live cell cross-correlation experiments corroborated the results obtained from lysates. IMPs were widely present in mRNPs, followed by ELAVL2 and STAU1. In the case of FMRP we observed a prominent cross-correlation between GFP-FMRP and mCherry-YBX1 pairs in live cells, indicating that the two factors diffuse together in live cells but cell lysis disrupts a putative RNA-independent association.

**Figure 7.**
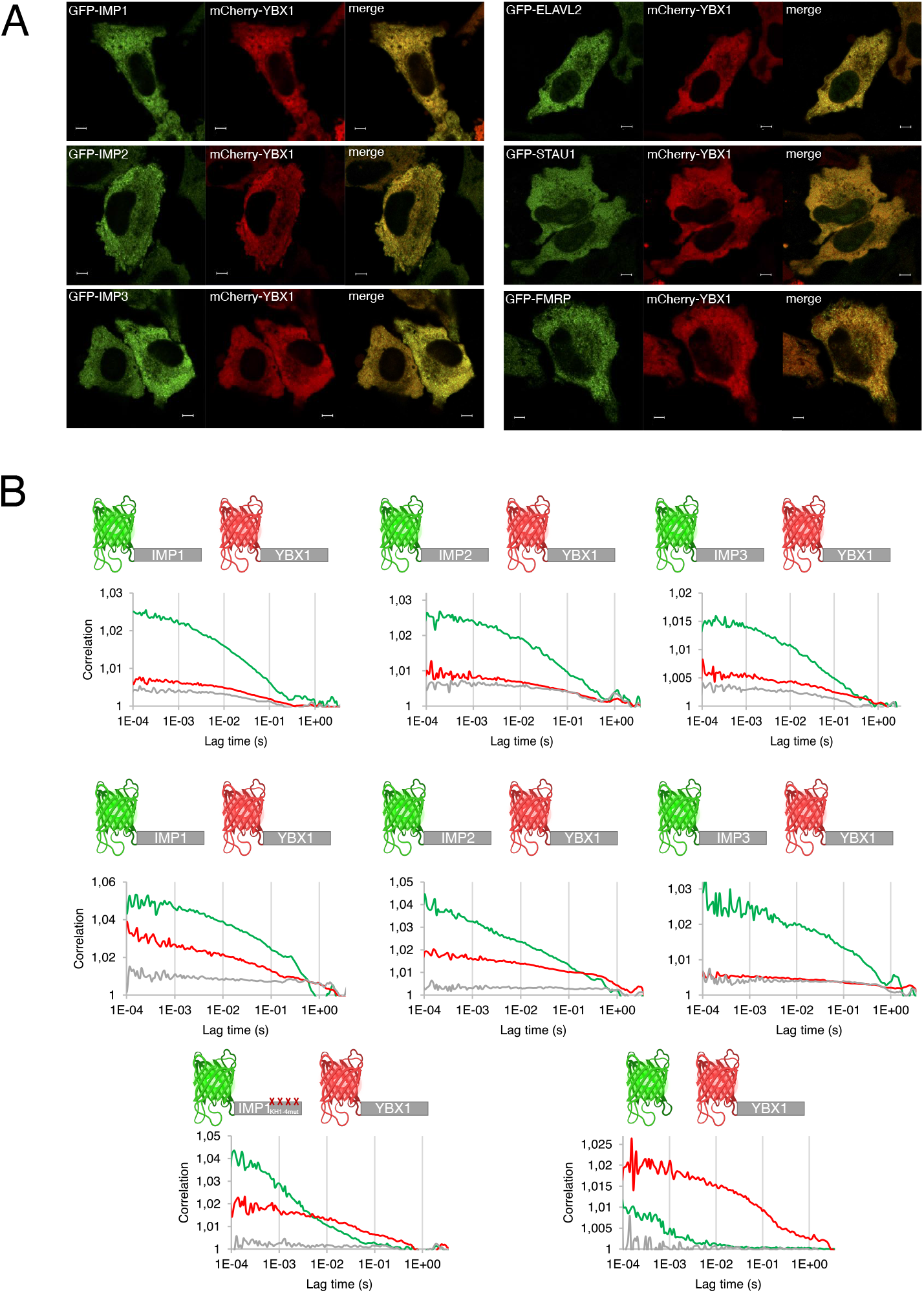
Protein-protein interaction quantification in live cells between mCherry-YBX1 and GFP-tagged IMP3, IMP2, ELAVL2, STAU1 and FMRP proteins. **A.** Confocal microscopy images of HeLa cells co-transfected with GFP-IMP1, GFP-IMP2, GFP-IMP3, GFP-ELAVL2, GFP-STAU1 or GFP-FMRP and mCherry-YBX1. Scale bar = 5 μm. **B.** Cytoplasmic live cell Fluorescence Cross-correlation Spectroscopy measurements of HeLa cells co-transfected with mCherry-YBX1 and GFP-IMP1, GFP-IMP2, GFP-IMP3, GFP-ELAVL2, GFP-STAU1 or GFP-FMRP. Co-transfection of mCherry-YBX1 and GFP-IMP1_KH1-4mut_ or GFP were used as negative controls for cross-correlation (no interaction).

## DISCUSSION

FCS is a broadly employable and versatile technology that has been applied to the molecular characterisation of fundamental biological processes such as plasma membrane organization, cellular transport, morphogenic gradients and chromatin organization in intact cells and organisms (22). FCS and FCCS per se do not provide direct structural information, as these techniques are based on a single-point measurement. This stands in direct contrast to image-based methods, which in turn do not resolve molecular interactions directly. FCS and FCCS have single molecule sensitivity and allow the analysis of molecular interactions under physiological conditions. Moreover, the procedures are quantitative and provide information about concentration, diffusion and interactions of particular proteins or larger protein complexes. From a practical perspective, FCS and FCCS are also appreciated as fast and highly reproducible methods that can be applied to studies of a wide range of fluorescent molecules. FCS and FCCS are appealing approaches to be used in the context of post transcriptional processes, governed by a series of combinatorial assemblies of proteins on RNA from transcription to mRNA translation and decay.

We applied FCS to both live cells and lysates because the two approaches complement each other and each have particular assets. Live cells can portray subcellular trafficking, docking or signalling events, whereas lysates have less spatial constraints and provide optimal readings of diffusion, stoichiometry and protein associations. Moreover, particular conditions or enzymatic treatments can be employed to lysates. Cytoplasmic mRNPs are stable entities, and factors such as IMPs remain attached to high affinity target mRNAs for hours (23). Consequently, lysates corroborate the results obtained in live cells, although diffusion is in general faster. Optimally - recordings should involve low concentrations of the fluorescent analyte (typically in the nM to μM range). In lysates this is adjusted by diluting the sample. In live cells, measurements should be performed in cells with low expression levels, and this can be readily identified in the accompanying confocal images which are typically taken with the same instrument prior to FCS data acquisition at a specific location of the cell. A limitation of FCS is obviously the need for the attachment of a fluorescent molecule which potentially can affect the function of the protein of interest. RNA-binding proteins such as IMPs and PABPC1 are sensitive to the position of tagged proteins and we experienced that separation of the fluorescent tag from the protein of interest by the addition of a flexible linker composed of Gly and Ser residues i.e. (GGGGS)_n_ (24) preserved the activity of the proteins - in particular YBX1.

Although CLIP- and RIP-seq analyses have provided a global picture of RBP attachments sites, IMP1 and YBX1 frequently exhibit multiple overlapping and mutually exclusive binding elements making it difficult to predict the exact composition of the individual mRNP (2,25,26). In the recorded fluorescence intensity fluctuations the mRNPs are readily visible as bright peaks as a consequence of oligomerization of RBPs to mRNA. The monomeric brightness can be obtained by the usage of an RNA binding mutant, cleavage of mRNA by RNase A or lysis in hypertonic buffer, allowing quantification of the number of RBPs per mRNP. This approach may essentially be applied to any RNA binding protein. As an example, we show that mRNPs on average contain 10 IMP molecules, which is roughly in agreement with the numbers of binding sites identified by CLIP (2,25).

Although changes in the diffusion coefficient of a protein may directly serve as an indication of its interaction with another protein-or molecular complex, FCS does not identify the binding partners of the protein. Consequently, the preferred method for detection of molecular interaction is FCCS. Compared to immunoprecipitation assays that typically require 10^7^-10^8^ million cells, FCCS requires only a single live cell, or about 10^4^-10^6^ cells if performed in lysates. We first examined the interactions of IMP1, YBX1 and PAPBC1. As expected, there was a strong cross-correlation between IMP1 and YBX1. The vast majority of IMP1 positive particles contain YBX1 and the same was observed with the two other IMP family members IMP2 and IMP3. While ELAVL2 showed a strong cross-correlation with YBX1 (similar to IMPs), cross-correlation was significantly lower with IMP1, indicating that IMP1 and ELAVL2 share the binding to some mRNAs but also bind to different mRNA subsets, uncovering a particular mRNP heterogeneity with these factors. In contrast, only a small proportion of STAU1 and FMRP were bound to YBX1 and IMP1 mRNPs in a RNA-dependent manner.

In conclusion, we report that FCS and FCCS are useful and reliable instruments for the analysis of mRNPs. Cytoplasmic mRNPs are in many ways suitable for FCS and FCCS analysis, but in principle we see no hindrance for the analysis of mRNPs in the nucleus or in defined domains such as synaptic termini where remodelling may occur. Moreover, FCS is moreover suitable for the study of assemblies that otherwise cannot be examined by methods relying on physical separation. Finally, we consider it an important feature of FCS that it produces quantitative data, providing a deeper understanding of the dynamic and heterogeneous assemblies.

## MATERIALS AND METHODS

### Vectors

Coding sequences of IMP1, IMP2, IMP3, ELAVL2, STAU1 and FMRP were PCR-amplified and cloned into pEGFP-C1 by restriction enzyme digestion. PABPC1 coding sequence was cloned into pEGFP-N1 using the same procedure. YBX1 was cloned into pcDNA3.1+N-eGFP inserting a 25 aminoacid flexible linker (5x GGGGS) between the fluorescent tag and YBX1. IMP1 and YBX1 (including flexible linker) coding sequences were also cloned into pmCherry-C1. EGFP was PCR-amplified and cloned into mCherry-C1 by restriction enzyme digestion in order to obtain mCherry-GFP fusion protein.

### Cell lines

HeLa cells (ATCC^®^ CCL-2^TM^) were grown in phenol red-free Dulbecco's Modified Eagle Medium (DMEM), high glucose, supplemented with GlutaMAX^TM^, 1 mM sodium pyruvate, 10% FBS (Biowest) and penicillin/streptomycin. A Flp-In^TM^ T-REx^TM^ 293 cell line (Invitrogen) stably carrying a 3xFLAG-IMP1 under a tetracycline inducible CMV promoter was generated and is referred in the text as TREX 293 3xFLAG-IMP1. TREX 293 3xFLAG-IMP1 cells were cultured in Dulbecco’s Modified Eagle Medium (DMEM), high glucose, supplemented with GlutaMAX^TM^, 10% FBS (Biowest), 5 µg/ml Blasticidin, 100 µg/ml Hygromycin and penicillin/streptomycin. Cell lines were cultured in a humidified incubator at 37ºC and 5% CO_2_.

### Plasmid transfections

400.000 HeLa cells were seeded in Nunc^TM^ cell-culture treated 6-well dishes. After 4-5 hours, cells were transfected with the plasmids described above in “Vectors” section using FuGene 6 and following the manufacturer’s instructions. Per dish, 1,3 μg of plasmid DNA and 3 μl of FuGene 6 (Promega) were mixed in 100 µl Opti-MEM^TM^ medium before the addition to the dish. For co-transfections, pEGFP and pmCherry vectors were mixed at 2:1 ratio.

### Confocal microscopy images

HeLa cells were seeded in 35 mm glass bottom dishes (No. 1.5 Coverslip, uncoated, MatTek), transfected with GFP or GFP/mCherry vectors and subsequently imaged ~24 hours after transfection, cells were fixed with 3,7% formaldehyde in PBS, washed and mounted in non-hardening VECTASHIELD antifade mounting medium (H-1000) with a refractive index n = 1.45. Confocal images were obtained using a Zeiss LSM780 confocal microscope with a Plan-Apochromat 63x/1.4 Oil objective.

### Fluorescence Correlation Spectroscopy (FCS)

18-22 hours post-transfection, cells were lysed at room temperature in 500 μl of lysis buffer containing 20 mM Tris-HCl pH 7.5, 140 mM KCl, 1,5 mM MgCl_2_, 1 mM DTT and 0,5% NP-40 supplemented with 1:300 mammalian protease inhibitor cocktail (Sigma). Cell lysates were briefly centrifuged at 800 xg for 1 minute and supernatant was transferred to a 35 mm glass bottom dish (No. 1.5 Coverslip, uncoated, MatTek) before being subjected to FCS. When indicated, cell lysates were treated with 100 ug/ml of RNase A (DNase and protease-free, EN0531, Thermo Scientific) and FCS measurements were recorded 1-2 minutes after RNase A treatment. For IMP stoichiometric measurements, cells were also lysed in 500 μl of hyperosmolar lysis buffer containing 20 mM Tris-HCl pH 7.5, 500 mM KCl, 1,5 mM MgCl_2_, 1 mM DTT and 0,5% NP-40 supplemented with 1:300 mammalian protease inhibitor cocktail (Sigma). FCS measurements were performed using a Zeiss LSM780 confocal microscope using a C-Apochromat 40x/1.2 W Corr M27 objective and using a water-phase immersion oil Immersol W 2010 (Zeiss). GFP measurements were performed with an argon laser with a 488 excitation wavelength and with a detection window between 500 and 633 nm (Supplementary Figure 1). Before each measurement, total count rate (kHz/s) was checked at different laser powers to ensure that fluorescence count signal was on the linear range. Live cell and cell lysate fluorescence measurements were recorded during 60 seconds and experimental autocorrelation curves, total count rate (kHz) and counts per particle (kHz) were obtained and analysed in ZEN 2011 software (Zeiss). Experimental autocorrelation curve fitting to 1-component and 2-component models was performed in ZEN 2011 software (Zeiss).

### Fluorescence Cross-correlation Spectroscopy (FCCS)

FCCS measurements were obtained after 18-22 hours of co-transfection with pEGFP- and mCherry-fusion protein plasmids. For live cell measurements, cells were located under the microscope and measurements were taken in arbitrary places in the cytoplasm. For cell lysate measurements, cells were lysed in 300 μl of lysis buffer containing 20 mM Tris-HCl pH 7.5, 140 mM KCl, 1,5 mM MgCl_2_, 1 mM DTT and 0,5% NP-40 supplemented with 1:300 mammalian protease inhibitor cocktail (Sigma). Cell lysates were subsequently subjected to FCCS and treated with 100 μg/ml RNase A (DNase and protease-free, EN0531, Thermo Scientific) when indicated. All FCCS measurements (live cell and cell lysate) were performed in a Zeiss LSM780 confocal microscope using a C-Apochromat 40x/1.2 W Corr M27 objective and Immersion oil Immersol W 2010 (Zeiss). GFP or mCherry measurements were performed with an argon laser with a 488 excitation wavelength and a DPSS laser with a 561 excitation wavelength, respectively. GFP fluorescence was captured with a detection window of 482-553 nm and mCherry fluorescence was captured with a detection window of 590-695 nm (Supplementary Figure 1). Before each measurement, total count rate (kHz/s) was checked at different laser powers to ensure that fluorescence count signal was on the linear range. Autocorrelation curves from live cell FCCS measurements shown in figures were recorded in arbitrary places in the cytoplasm during 20 seconds (live cell FCCS) or 60 seconds in the case of cell lysate FCCS. Experimental autocorrelation and cross-correlation curves were obtained and analysed in ZEN 2011 software (Zeiss).

### 3xFLAG-IMP1 immunoprecipitation

TREX 293 3xFLAG-IMP1 cell line was used for immunoprecipitation. 5 × 10^6^ cells were seeded in 140 mm diam. Nunc^®^ petri dishes and expression of 3xFLAG-IMP1 was induced by the addition of 1 µg/ml tetracycline. Cells were transfected 4-5 hours after seeding using FuGene 6 following the manufacturer’s instructions with pEGFP plasmids described in “Vectors” section. Per dish, 16.75 μg of plasmid DNA and 37.5 μl of FuGene 6 were mixed in 1200 μl Opti-MEM^TM^ medium before addition to the cells. 48 hours after seeding, cells were lysed in 1 ml of lysis buffer containing 20 mM Tris-HCl pH 7.5, 140 mM KCl, 1.5 mM MgCl2, 1 mM DTT and 0.5% NP-40 supplemented with 1:300 mammalian protease inhibitor cocktail (Sigma) on ice. Cell lysates were cleared at 8200 xg for 5 minutes at 4ºC. 50 µl of Anti-FLAG^®^ M2 Magnetic beads (Sigma) were added to each cleared cell lysate and samples were incubated for 3 hours with rotation at 4ºC. Beads were washed 3x with lysis buffer and finally immunoprecipitated material was collected by addition of 2x SDS buffer directly to the beads.

### Western Blot

Total lysate and FLAG immunoprecipitation samples were loaded into a 10% RunBlue^TM^ SDS protein gel (Expedeon) and separated by electrophoresis. Proteins were transferred to a PVDF membrane using the iBlot^TM^ 2 Gel Transfer system (Invitrogen). After transfer, membrane was blocked in wash buffer with 5% skim milk powder for 1 hour at room temperature. After blocking, primary antibodies against GFP (rabbit mAb, Cell Signaling) and FLAG (M2 mouse mAb, Sigma) were incubated overnight at 4ºC. The day after, membranes were washed and incubated with either anti-rabbit or anti-mouse HRP-conjugated secondary antibodies (Cell Signaling). After incubation, membranes were washed, incubated 2-3 minutes with SuperSignal^TM^ West Pico Chemiluminiscent Substrate (Thermo Scientific) and imaged using C-DiGit^®^ Blot Scanner (LI-COR).

## Acknowledgements

The authors would like to thank Stine Østergaard for her technical assistance.

## Author contributions

À.M-R. and F.C.N. designed the study. À.M-R. performed the experiments. À.M-R. and F.C.N analysed the data. À.M-R., J.C., C.H. and F.C.N. wrote the manuscript.

