## Supplementary Material for "Fluorescence Correlation and Cross-Correlation Spectroscopy Unveil Cytoplasmic mRNP Composition and Dynamics"

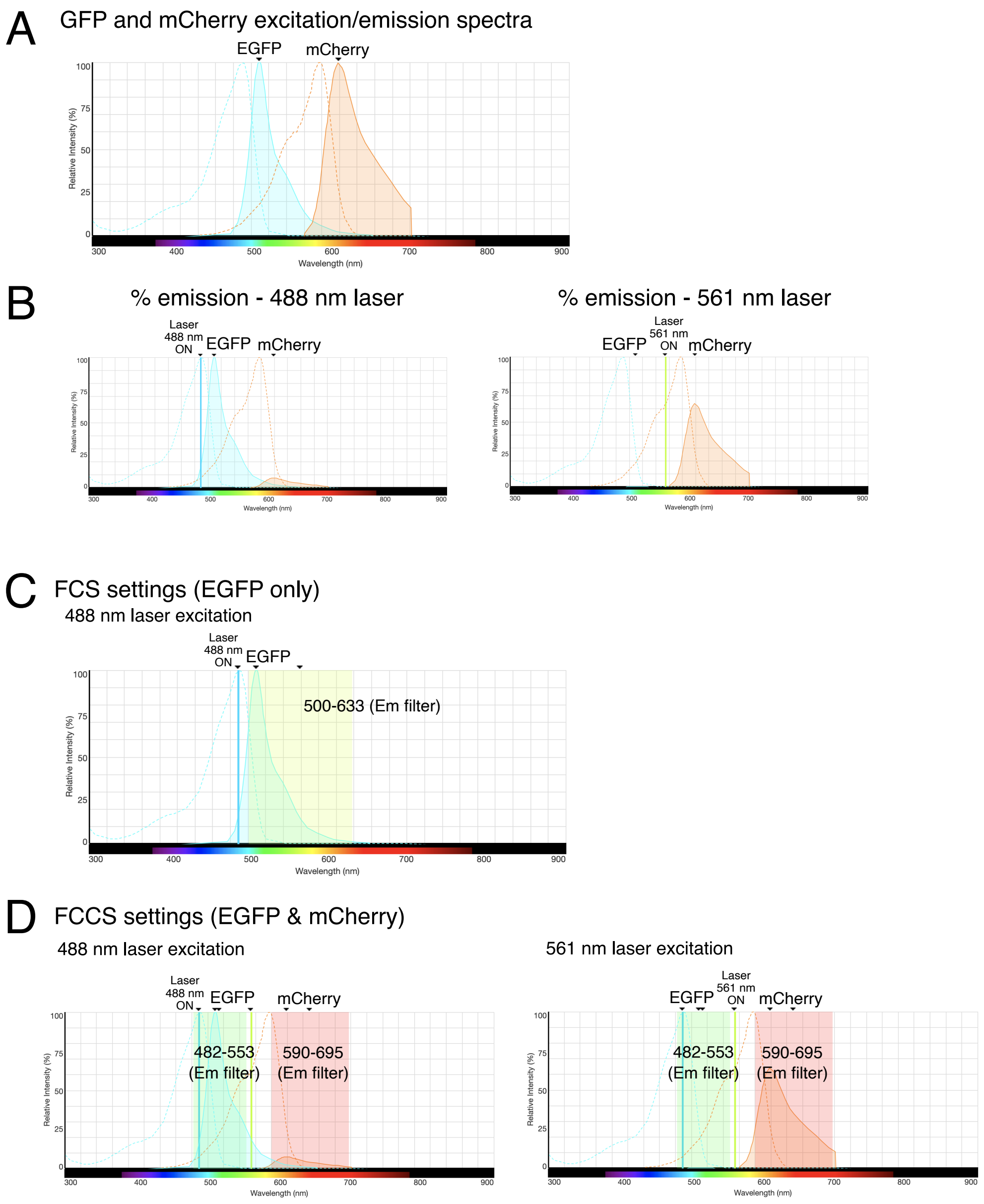


**Supplementary Figure 1. EGFP and mCherry emission/spectra and FCS/FCCS settings.** **A.** GFP (blue) and mCherry (orange) excitation and emission spectra. Discontinued curve represents emission spectra and filled curve represents emission spectra. **B.** Relative intensity (%) of emission of GFP and mCherry upon excitation with 488 (left) or 561 (right) nm lasers. **C.** Representation of FCS excitation laser wavelength and emission filter settings for GFP only FCS measurements. **D.** Representation of FCCS excitation laser wavelength and emission filters for GFP and mCherry FCCS measurements. Relative intensity (%) or GFP and mCherry upon 488 (left) or 561 (right) nm laser excitation.

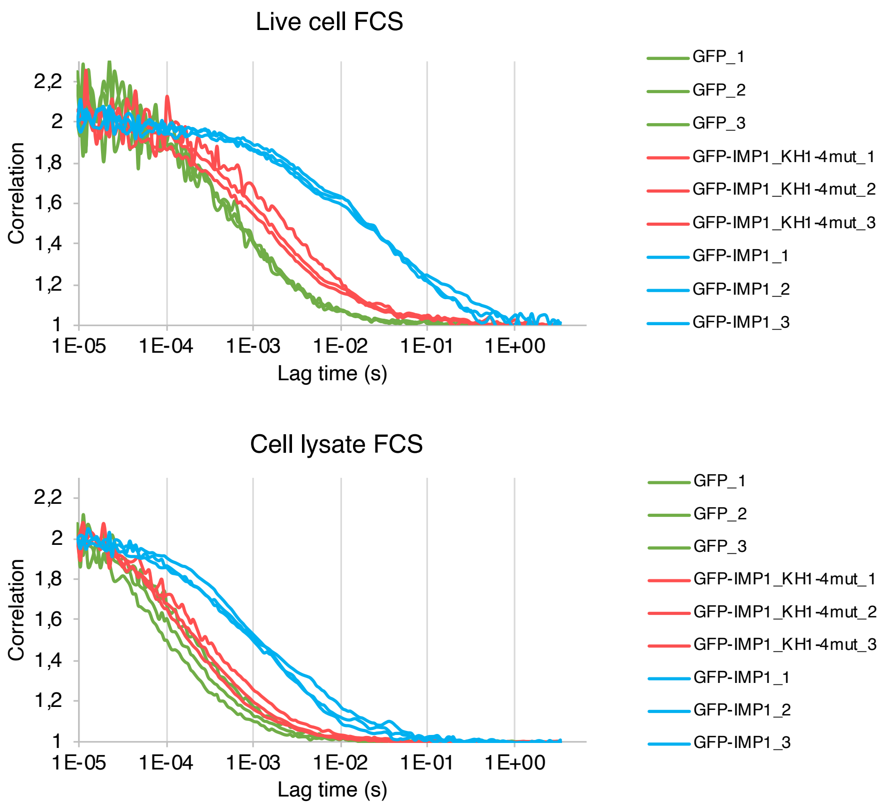


**Supplementary Figure 2. Inter-assay variation of FCS measurements in live cells and cell lysates.** Normalized autocorrelation curves from three biological replicates of cells transfected with GFP, GFP-IMP1$_{KH1\text{-}4mut}$ and GFP-IMP1.

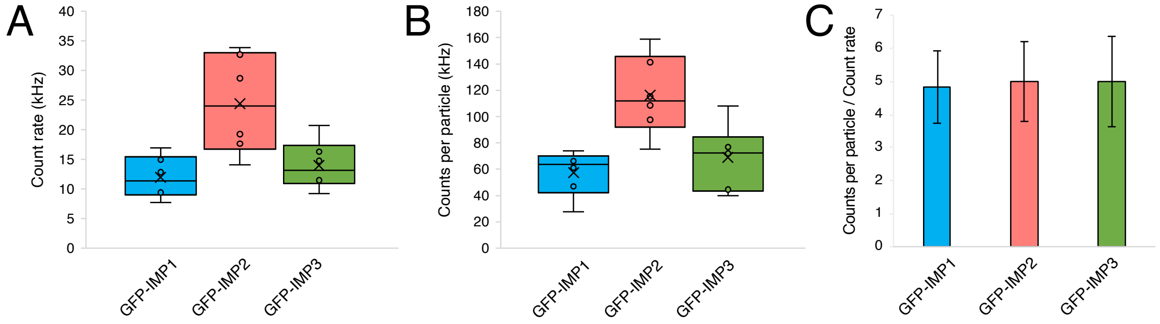


**Supplementary Figure 3. A.** Total count rate (kHz/s) measurements in cell lysates of cells transfected with GFP-IMP1, GFP-IMP2 and GFP-IMP3 (n=6 biological replicates). **B.** Correspondent counts per particle (kHz) measurements in cell lysates of cells transfected with GFP-IMP1, GFP-IMP2 and GFP-IMP3 (n=6 biological replicates). **C.** Counts per particle normalized against total count rate.

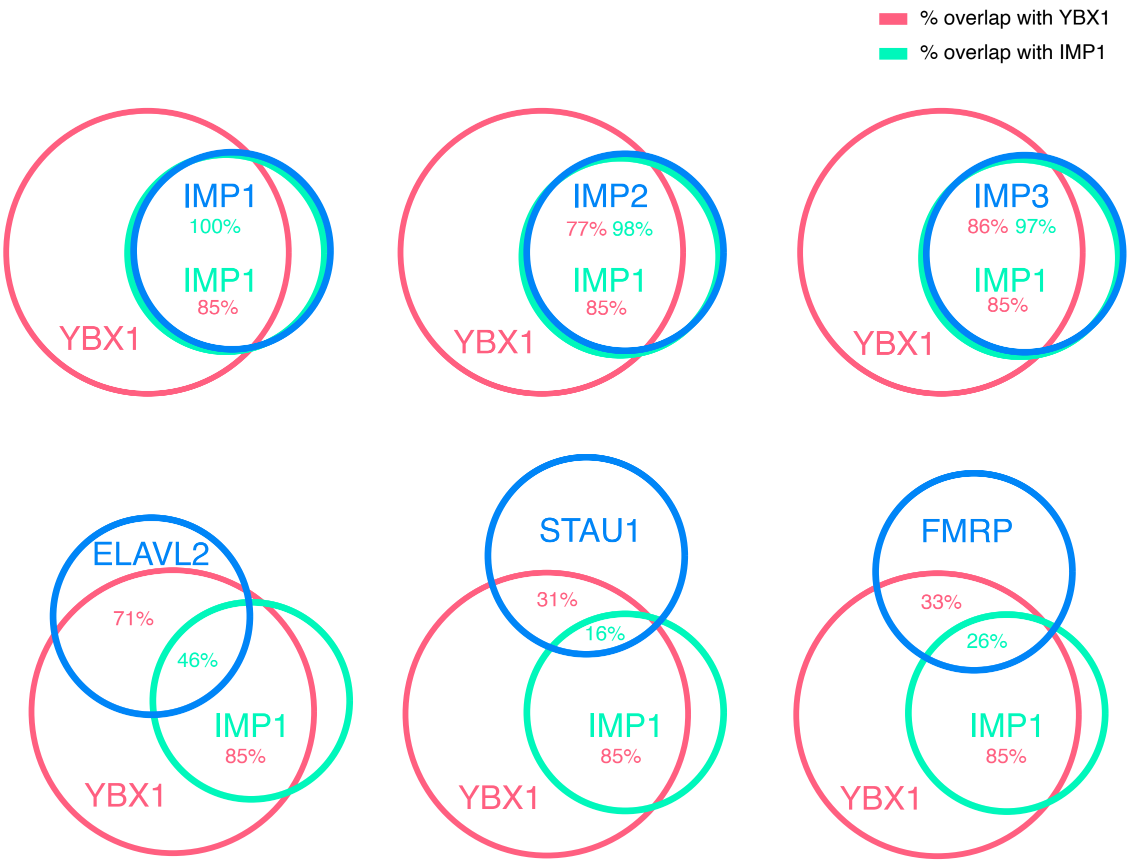


**Supplementary Figure 4.** Model of mRNP heterogeneity showing all the factors tested and its correspondent % overlap with YBX1 and IMP1 according to FCCS results. Average % are obtained from cross-correlation ratios shown in Figure 5 and Figure 6. A cross-correlation of 0.7, obtained in the GFP-IMP1 and mCherry-IMP1 FCCS measurement is considered to be 100% and all the % shown are calculated accordingly.
